# Sensing hand position in Ehlers-Danlos Syndrome

**DOI:** 10.1101/2021.04.09.439251

**Authors:** Holly A. Clayton, Bernard Marius ’t Hart, Denise Y. P. Henriques

## Abstract

**Purpose:** To explore the effect of joint hypermobility on acuity, and plasticity, of hand proprioception.

**Materials and Methods:** We compared proprioceptive acuity between EDS patients and controls. We then measured any changes in their estimate of hand position after participants adapted their reaches in response to altered visual feedback of their hand. The Beighton Scale was used to quantify the magnitude of joint hypermobility.

**Results:** There were no differences between the groups in the accuracy of estimates of hand location, nor in the visually-induced changes in hand location. However, EDS patients’ estimates were less precise when based purely on proprioception and could be partially predicted by Beighton score.

**Conclusions:** EDS patients are less precise at estimating their hand’s location when only afferent information is available, but the presence of efferent signalling may reduce this imprecision. Those who are more hypermobile are more likely to be imprecise.

## Introduction

Ehlers-Danlos syndrome (EDS) is a group of genetic connective tissue disorders that can afflict up to 2% of individuals. Most forms of EDS affect collagen throughout the body; some directly impact its structure (such as with Classical EDS), while others alter proteins that interact with collagen [1]. Although symptoms can vary across, or sometimes within, each of the sub-types, the feature that most EDS patients have in common is joint hypermobility (see [2] for a more detailed review of variations in genetics, and symptomology, across the thirteen subtypes).

Motor control requires knowledge of where our limbs are in space. Impaired sense of proprioception can sometimes lead to movements that appear ‘clumsy’. EDS patients are frequently described as exhibiting clumsy movements, for which impaired proprioception is offered as an explanation [3]. Indeed, a limited number of studies have suggested that EDS patients, or other groups exhibiting joint hypermobility, may have proprioceptive impairments [4–8], but the exact nature of this impairment is still unclear. Specifically, we are not sure whether this impairment only emerges during passive proprioception (which should only rely on afferent information), or whether it also occurs during active proprioception (which should Sensing hand position in EDS rely on both afferent and efferent information). In addition to furthering our understanding of proprioceptive acuity in EDS, we also test the extent to which proprioception is affected by visual misalignment of the hand during reaching movements. The goal of this study is to understand how joint hypermobility, which is typically seen in those with EDS, affects both the accuracy (average) and precision (variability) of estimates of hand position both before and after sensorimotor adaptation.

Previous research from our lab suggests that EDS patients do show differences in proprioceptive sensitivity. Most recently we found that, although patients were just as accurate as controls, they were significantly less precise when indicating the felt position of their left hand at 6 different locations in a horizontal workspace. Specifically, patients showed twice as much scatter in these judgements compared to controls at all locations [4]; this suggests proprioception is less precise in EDS. The greater amount of scatter did not correlate with the magnitude of chronic pain, suggesting that pain was not contributing to the proprioceptive deficit here. In another study of ours [5], we again found that EDS patients showed proprioceptive estimates that were of similar accuracy as controls, and the precision of these estimates was significantly worse (around half of that of controls), but only at locations lateral to the body midline. The precision of these estimates at peripheral locations was significantly correlated with Beighton scores, which are commonly used to measure the magnitude of joint hypermobility. In other words, we found that those who were the most hypermobile were also the least precise when estimating their hand at peripheral locations. This suggests that hypermobility could be related to the proprioceptive deficit that seems to occur in EDS.

However, in both studies mentioned above, for proprioceptive assessment, participants moved their own unseen hand along robot-generated slots to their final location. Thus, the estimates of their unseen hand may not have been purely based on proprioceptive information since the participant had to push their hand to the final site. Yet their hand path was constrained, and its direction and final location varied across trials, so they also could not benefit from the extra information contained in efferent signals that would have been fully available if the hand direction had been entirely generated by the participant themselves. It is possible that proprioceptive differences would have been even larger in EDS participants if their hand had been passively carried to its final location. Therefore, we want to know the extent that additional efferent information (produced during self-generated movements) can attenuate these proprioceptive deficits. To test for this, we measured both (1) estimates of hand location after the hand was passively displaced, using a robotic manipulandum and (2) estimates of hand location after the hand was actively displaced, by the person themselves, at a self-chosen location. We compared proprioceptive acuity in both tasks between EDS patients and controls. Based on our past research [4–5], we hypothesized that EDS patients’ proprioceptive localizations would be similar in accuracy to those of healthy controls, but that patients’ localizations would be significantly less precise. We also anticipated that having efferent information available would reduce the expected imprecision in EDS.

Our second goal was to measure proprioceptive plasticity in EDS. For this goal, we altered visual feedback of the hand during a reach-training task, and afterwards measured how training with this visual distortion shifted estimates of hand location. Again, we compared the extent of the visual-induced changes between EDS patients and controls, and further, whether Beighton scores were related with proprioceptive acuity or plasticity. We hypothesized that EDS patients would adapt their reaches to a similar extent as healthy controls, that their estimates of hand location would shift similarly and that we would find a significant correlation between our measures of hand proprioception and Beighton scores, like in our first study [5]. Our results confirm that proprioceptive information is less precise in EDS, but may be slightly attenuated by efferent information, and that proprioceptive precision is partially related to the magnitude of joint hypermobility. Our results provide a more comprehensive understanding of the proprioceptive sensitivity in EDS.

## Materials and Methods

### Participants

Sixteen healthy controls (mean age 34 years, range 18-54, 13 females) and fourteen EDS patients (mean age 34 years, range 25-37, 11 females; 3 Classical EDS, 9 Hypermobile EDS, 1 Arthrochalasia EDS and 1 Spondylodysplastic EDS) voluntarily took part in the experiment outlined below. All participants had corrected-to-normal vision and were right-handed. Controls were laboratory volunteers or recruited from the Undergraduate Research Participant Pool at York University (and given course credit for their participation). Participants in the patient group were recruited through EDS Canada’s General Toronto Area Support Group. All participants provided informed consent, and the study was conducted in accordance with the ethical guidelines set by the York Human Participants Review Subcommittee.

EDS was recently re-classified into thirteen sub-types [2], after identifying the genetic mutations responsible for twelve of the sub-types (genetics responsible for Hypermobility type are still unknown). Only patients with confirmed diagnoses (confirmed clinical diagnoses for Hypermobility type; confirmed molecular diagnoses for all other types) were admitted into the study. Joint hypermobility was measured using the Beighton criterion, which rates patients’ hypermobility on a 9-point scale after performing 9 movements. Patients’ Beighton scores were obtained from genetic reports and were confirmed by the experimenter prior to testing with a goniometer. None of the EDS patients were on any medication known to affect their cognitive abilities during the experiment.

### General Experimental Setup

Participants sat on a height and distance adjustable chair in front of the experimental set-up. With their right hand, participants held onto the vertical handle of a two-joint robot manipulandum (Interactive Motion Technologies Inc., Cambridge, MA, USA) such that their thumb rested on top of the handle. A black cloth was draped over their shoulder, and right arm, to occlude visual feedback of the reaching limb. Visual stimuli were projected from a downward facing monitor (Samsung 510 N, 60 Hz) located 28 cm above the robotic arm. A reflective surface was mounted on a horizontal plane 14 cm above the two-joint robotic arm, midway between the manipulandum and the monitor, such that images displayed on the monitor appeared to lie in the same horizontal plane as that of the robotic arm (figure 1A). Underneath the reflective surface, ~2 cm above the position of the thumb, as it rested on the modified handle of the manipulandum, a touch screen was mounted so participants could indicate unseen right-hand locations (specifically the unseen thumb) with their left hand, for some tasks. The left hand was illuminated by a small lamp during these tasks, and therefore was visible when reaching to the touch screen panel. For each task, the home position of the right hand was located ~20 cm in front of the participants, along the participants’ body midline.

**Figure 1.**
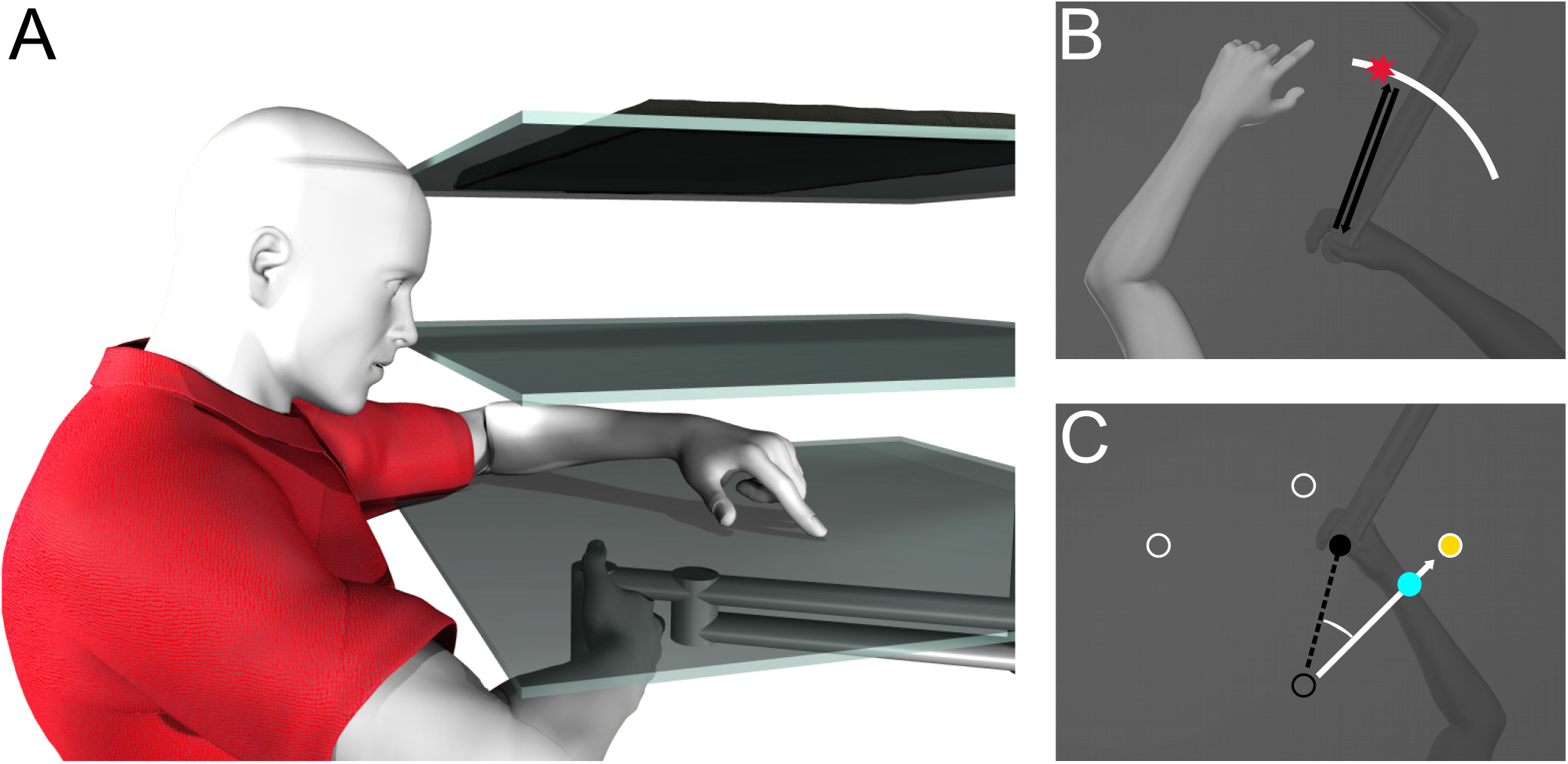
Setup and experimental design. **A:** Participants moved their right hand which was hidden by a mirror (middle surface) half-way between their hand and the monitor (top surface). A touchscreen located just above the right hand was used to collect responses from the left hand (bottom surface). **B:** Active and Passive Localization Trials. One of three white arcs, spanning 60°, located 12 cm away from the home position and centred at 50°, 90° or 130° would appear. Only the 50° arc is shown as an example. The participants’ invisible, right hand first moves to the arc and back to the home position, either by their own voluntary movement (“active localization”) or by the robot (“passive localization”), indicated by the black pathway. Then they use their visible, left hand to indicate on the touch screen where their right hand crossed the arc, indicated by the red star. The home position is not shown to prevent it from being used as a reference point (the hand is at the home position in the illustration and is at the same position as the open black circle in panel C). **C:** Reach training task and No-Cursor reach trials. The targets were located 12 cm away from the home position (shown by the hollow black circle) at 45° (shown here by the yellow circle), 90°, and 135° (shown by the hollow white circles) and were presented one at a time in a pseudorandom order. In the rotated training tasks, the hand-cursor (blue circle) was rotated 30° relative to the home position. In the aligned training task, the cursor was green and aligned with the hand’s position (not depicted here). In the No-Cursor trials, the cursor was not visible, thus no visual feedback of the hand’s position was available.

### Procedure

All participants completed the set of tasks in a specified order in two sessions performed one after the other (see figure 2). Each session started with a reach training task, followed by several localization tasks, as detailed below. In between the localization tasks there were blocks of no-cursor reaches and additional training.

**Figure 2.**
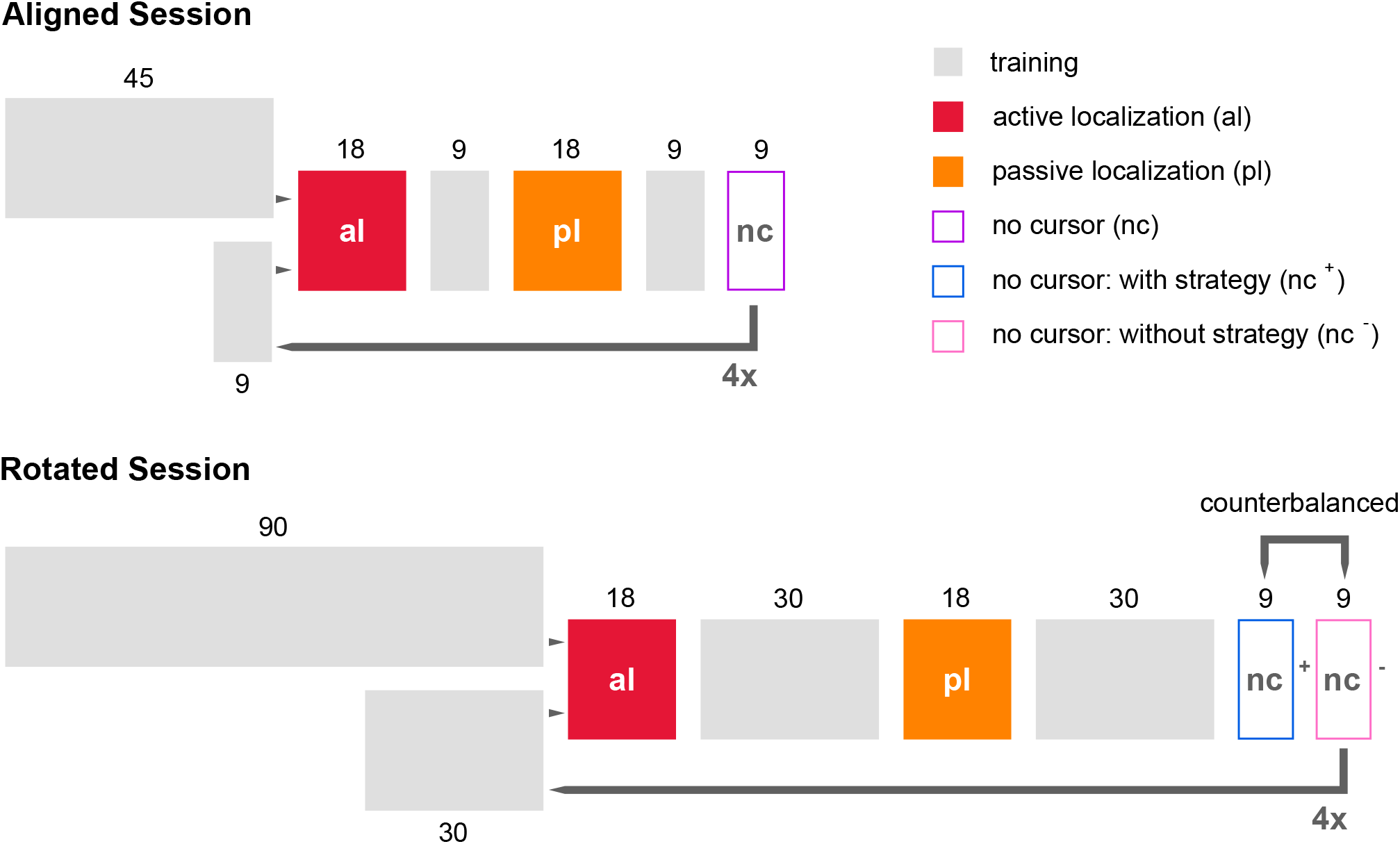
Experiment paradigm detailing the order of tasks, and number of trials, across aligned and rotated sessions. **Top:** For the first session the cursor was aligned with the position of the right hand. Participants began with 45 cursor training trials that were then followed by blocks of active localization (red, 18 trials each), passive localization (orange, 18 trials each) and no-cursor trials (hollow, 9 trials each). Nine ‘top up’ aligned-cursor training blocks were interleaved in between localization and no-cursor blocks for four more repeats. **Bottom:** During the second session the cursor was rotated 30° CW relative to the position of the right hand. Participants began with 90 cursor training trials that were then followed by blocks of active localization (18 trials each), passive localization (18 trials each) and two variations of blocks of no-cursor trials (with or without strategy; 9 trials each). Each block was followed by 30 ‘Top up’ rotated-cursor training blocks for four more repeats. In both the aligned and rotated sessions, passive localization always occurred after active localization, since endpoint locations of the robot-generated movements in passive localizations were based on locations that participants voluntarily moved towards during active localization.

The aligned session consisted of four blocks of reach training, hand localization and no-cursor trials; this served as baseline data (Aligned session; figure 2). The session began with 45 aligned (green) cursor training trials, followed by blocks of 18 active localization trials, 18 passive localization trials, and 9 no-cursor trials. Shorter blocks of 9 cursor training trials, referred to as ‘top-up’ cursor training trials, were interleaved between localization and no-cursor blocks.

Following the aligned session, participants were told that the cursor would be moving differently, and that they would have to compensate for the difference, but they were not explicitly informed that the cursor’s trajectory would be rotated 30° CW from their actual hand’s movement. During the rotated session, the cursor was blue, and its motion was rotated 30° CW relative to the home position. To perfectly compensate for this visuomotor rotation, the unseen hand would have to move in a direction 30° CCW from any displayed target relative to the home position. The rotated session (figure 2) began with a longer training session of 90 rotated training trials and each top-up block contained 30 trials, to reduce learning decay of the visuomotor rotation. Each block of rotated reach training trials was followed by 18 trials of active localization, 18 trials of passive localization and 18 no-cursor trials. Each block of no-cursor trials was done twice, although only one set (those where participants were asked not to employ a strategy during reaching, see task description below) was used for this analysis.

### Hand Localization

The hand localization tasks (figure 1B) were used to measure acuity of unseen hand localization and have been established as a reliable measure of proprioceptive abilities [9–14]. In the active hand localization tasks, participants moved their own hand to a self-chosen position on the arc, and thus both afferent and efferent information was available. In the passive hand localization task, the robot displaced their passive hand, such that only afferent information on hand location was available. The robot displaced the hand to the same endpoints that were recorded in the preceding ‘active’ task (to ensure no differences in hand-target locations across conditions), but in a shuffled order, which required the active hand localization task to always be performed first.

Each hand localization trial began with a white arc (0.5 cm thick, located 12 cm away from the home position, which was not visible to participants because it could provide a reference point) appearing on the screen (figure 1B; home position is at the same location as the right invisible hand). The arc spanned 60° and was centred on either the 50°, 90° or 130° location in polar coordinates, and the target-hand was moved 12 cm out (either by the participant or by the robot) until the hand hit a force cushion. With the arc still displayed, participants used their visible left hand to indicate, on the touch screen mounted above the manipulandum, the location where the movement of their unseen right hand had crossed the arc (comparable to [9]; participants were to point with the left hand to ‘where they believed their right hand crossed the circle’). After each touchscreen response participants were instructed to place their left hand under their chin to prevent unintended contact with the touchscreen.

### Training

Besides measuring estimates of hand location, we also wanted to measure how these estimated locations change with visuomotor training [15]. Visuomotor adaptation is when people reach to targets with a misaligned cursor, representing their unseen hand, and is considered a reliable method for studying visuomotor learning [16]. Visuomotor training involved reaches to a single, visual target (a yellow disc with a diameter of 1 cm), 12 cm away at 45°, 90° or 135° relative to the home position (figure 1C). Participants were instructed to reach to the target as quickly and as accurately as possible using a green (aligned session) or blue (rotated session) circular cursor, 1 cm in diameter, representing their unseen hand. A reach trial was complete when the centre of the hand cursor overlapped with the target (i.e. the hand was within 0.5 cm of the target’s centre). Upon completion of the reach, both the cursor and target vanished, and the participants moved their hand back toward the home position, along a constrained, straight path. That is, if participants tried to move outside of the path, a resistance force (a stiffness of 2 N/(mm/s) and a viscous damping of 5 N/(mm/s) was generated perpendicular to the path. During aligned-cursor training the cursor was aligned with movement of the unseen hand. During rotated-cursor training the motion of the cursor was abruptly rotated 30° clockwise relative to the home position where it remained for all subsequent trials and blocks.

### No-Cursor Reaches

Reach after-effects are measured by having participants reach to targets in the absence of the hand-cursor and are considered a reliable measure of implicit learning [16]. Participants reached to each of 3 targets: 45°, 90°, and 135°, three times each, pseudo-randomly, for a total of 9 reaches per block (figure 2). After the hand moved out and was held in the same position for 300 ms, the target disappeared which indicated that the trial was over. Participants then returned their hand to the home position along a constrained pathway, like in training.

During the rotated session, we used two variations of No-Cursor trials (including and excluding a strategy; figure 2). Participants completed these two variations in succession and the order was counterbalanced across participants. Participants were instructed to either include or exclude any strategy that they may have developed to counter the visuomotor rotation during rotated cursor training trials, even though they were not explicitly told how to counter it. These tasks were inherited from other studies in our lab, involving healthy participants, which measured explicit and implicit processes of visuomotor adaptation [11–14]. Out of convenience we used the same set of programmed tasks for the current study, but since our goal here was to measure proprioceptive acuity in EDS (and we did not think implicit/explicit contributions would be relevant here) we planned a priori to exclude the “with-strategy” trials from analysis.

### Data Analysis

The main goal of this experiment was to determine the effect of EDS on the accuracy and precision of both active and passive hand localizations. To put any such effect in the proper context, we first tested if there were any differences in performance in visuomotor learning (table 1; first two tasks). Finally, we investigated the relation between hypermobility and hand localization. For all statistical tests, the alpha level was set to 0.05 and, when appropriate, Greenhouse-Geisser corrections were used. A summary of the measures derived from each task are presented in table 1 and are described further below. All data pre-processing and analyses were done in R version 3.6.0 (R Core Team, 2019). All data and analysis scripts are available on OSF (https://osf.io/t2jrs/).

**Table 1.** A summary of the measures that were used to analyse results from each of the experimental tasks.

| **Task** | **Accuracy Measure**<br>(Aligned or Rotated) | **Accuracy Difference Measure**<br>(Rotated - Aligned) | **Precision Measure**<br>(Aligned or Rotated) |
| --- | --- | --- | --- |
| *Training* | Mean angle of movement at maximum velocity; not shown in figures. | Reach deviations; Figure 3 (A-C). | Standard deviation of angle at maximum velocity; Figure 3D. |
| *No-cursor Reaching* | Mean angle at movement endpoint; Figure 4A. | Aftereffects; Figure 4B | Standard deviation of angle at movement endpoint; Figure 4 (C-D). |
| *Active Localization* | Smoothed-spline interpolation through the angular differences between the location where the participant's unseen, self-generated, right-hand movement ended and the perceived hand location, where participants indicated on the touchscreen with their left hand; Figure 5A (aligned only; rotated not shown). | Active shifts; Figure 5C | Mean squared error between the angular location of the perceived hand position and a smoothed-spline interpolated reference. This assumes the smoothed-spline as equivalent to a mean, to approximate the standard deviations in the other tasks; Figure 6. |
| *Passive Localization* | Smoothed-spline interpolation through the angular differences between the location where the participant's unseen, robot-generated, right-hand movement ended and the perceived hand location, where participants indicated on the touchscreen with their left hand; Figure 5B (aligned only; rotated not shown). | Passive shifts; Figure 5D | Mean squared error between the angular location of the perceived hand position and a smoothed-spline interpolated reference. This assumes the smoothed-spline as equivalent to a mean, to approximate the standard deviations in the other tasks; Figure 6. |

### Rate of Adaptation

First, we analysed group differences in rates of learning for reaches during cursor training trials (table 1; first task row). All cursor, and no-cursor reaches, included in both sessions (aligned and rotated) were manually inspected to ensure participants performed the task as requested. For example, a trial would be removed if a participant did not attempt to reach directly towards the target. The very small number of trials (less than 5% in each group) that were found to have violated the instructions were removed from further analyses. For the remaining trials, we calculated angular reach deviation at the point of maximum velocity. We corrected for individual baseline biases, by calculating the average reach deviation for each target separately within each participant, during the last 30 out of the first 45 aligned-cursor training trials. The first set consisted of the first 3 trials (trials 1-3), the second consisted of the next 3 trials (trials 4-6), and the third consisted of the last 15 trials of training (trials 76 to 90). We then compared measures of angular reach deviation, for each of these three trial set across patients and controls using a 3 × 2 mixed design ANOVA; this allowed us to confirm whether both groups learned to counter the perturbation and to explore any differences across both groups.

We also computed the standard deviation of both cursor and no-cursor reaches, for both the aligned and rotated sessions, as a measure of precision which we compared across both groups by using two separate 2 × 2 mixed design ANOVAs.

### Reach After-effects

Then we explored possible group differences in reaching movements when cursor feedback was absent (table 1; second task row). We took the angular reach deviations at movement endpoints for these no-cursor, open-loop, reaches. For all no-cursor trials, we calculated the angular difference between a straight line from the home position to the point where the participant’s hand movement ended, and a line from the home position to the target. Using the endpoint of the reach, rather than the point of maximum velocity, makes data more comparable to those obtained in localization trials (see below).

To measure implicit learning following training with a rotated cursor, angular reach deviations from aligned no-cursor trials were subtracted from without strategy no-cursor trials. Since we were only interested in implicit motor adaptation, we only looked at the reach aftereffects without strategy. We compared implicit learning across both groups by using a 2 × 2 mixed design ANOVA.

### Hand localization

To answer our main questions, we explored the effect of EDS on hand location estimates. To do so, we analysed hand location estimates (both after active and passive hand displacement) before and after visuomotor adaptation (table 1; last two task rows). We computed the angular difference between a line connecting the home position to the location where the participant’s unseen right-hand movement ended, and a line connecting the home position to perceived hand location (where participants indicated on the touchscreen with their left hand). To account for possible differences in performance of these localization tasks, we ensured that arc responses were centred where we expected the arc to be displayed, 12 cm from the home position, by using the same circle-fitting procedure as another study from our lab [12]. This helped to ensure that any localization shifts detected in analyses were not due to unwanted response biases or technical issues. Furthermore, all localizations included in both sessions were manually inspected to ensure participants performed the task as requested, just like we did for our reaching tasks, and only a few trials were excluded from analysis. Since participants chose the locations on the arc that they moved towards in the Active Localization task, their movements did not always encompass all possible arc locations. Thus, as we did in our other studies which used the same tasks [11–14], we incorporated a kernel-smoothing method (gaussian kernel with a width of 15°) to interpolate changes in hand localization at specific points (50°, 90° and 130°; the same locations where the arcs were centred in polar coordinates) for every participant. In order to confirm that baseline hand localizations were no different between the groups we calculated 95% confidence intervals, based on sample t-distributions, and plotted them for both groups, across each of the three hand angles, separately for each of the localization tasks (active and passive), as a way of testing for significant differences since this is equivalent to running a t-test but has the advantage of showing the distribution of the data. Then we used the mean of these values, at each of the three points, to estimate the accuracy of hand localization errors in active and passive movements for both the aligned and rotated sessions, which we compared across both groups using a 2 × 2 × 2 mixed design ANOVA. We also computed the standard deviation of the hand location estimates, for both afferent-based estimates and efferent-based estimates in both the aligned and rotated sessions, as a measure of proprioceptive precision which we compared across both groups using a 2 × 2 × 2 mixed design ANOVA. Then we conducted a series of 1-sided Welch t-tests to explore group differences across each of the 4 versions of this task.

To measure the effect of visuomotor adaptation on hand localization, we first confirmed that hand localization after rotated-cursor training significantly differed from that after aligned-cursor training as in our previous studies. Then, we calculated the difference of localization errors between the two sessions to represent visually-induced shifts in hand localization. These shifts were compared across groups, separately for active and passive hand localization, using two separate 2 × 2 mixed-design ANOVAs.

## Results

### Learning Rate

Before investigating how EDS affects changes in hand localization, we first confirmed that both groups appropriately countered the perturbation by the end of 90 training trials (figures 3A-C). We tested for group differences in reach deviations at different time points during adaptation training (three blocks: trials 1-3, 4-6, 76-90) using a 3 × 2 mixed design ANOVA, with block as a within-subject factor (blocks 1, 2 and 3) and group as a between-subject factor (control and EDS). We found a statistically significant effect of block (F (1.63, 45.72) = 41.117, p < .001, η^2^ = 0.459) and Tukey post-hoc analyses revealed that there were significant differences between all three of the blocks (figure 3B). However, we found no statistically significant differences between the two groups (F (1, 28) = 0.271, p = .777, η^2^ = 0.001), nor an interaction between group and block (F (2, 56) = 0.114, p = .892, η^2^ = 0.002). This suggests that, as expected, both groups learned and that there is no discernible difference in their rate or asymptotic level of learning.

**Figure 3.**
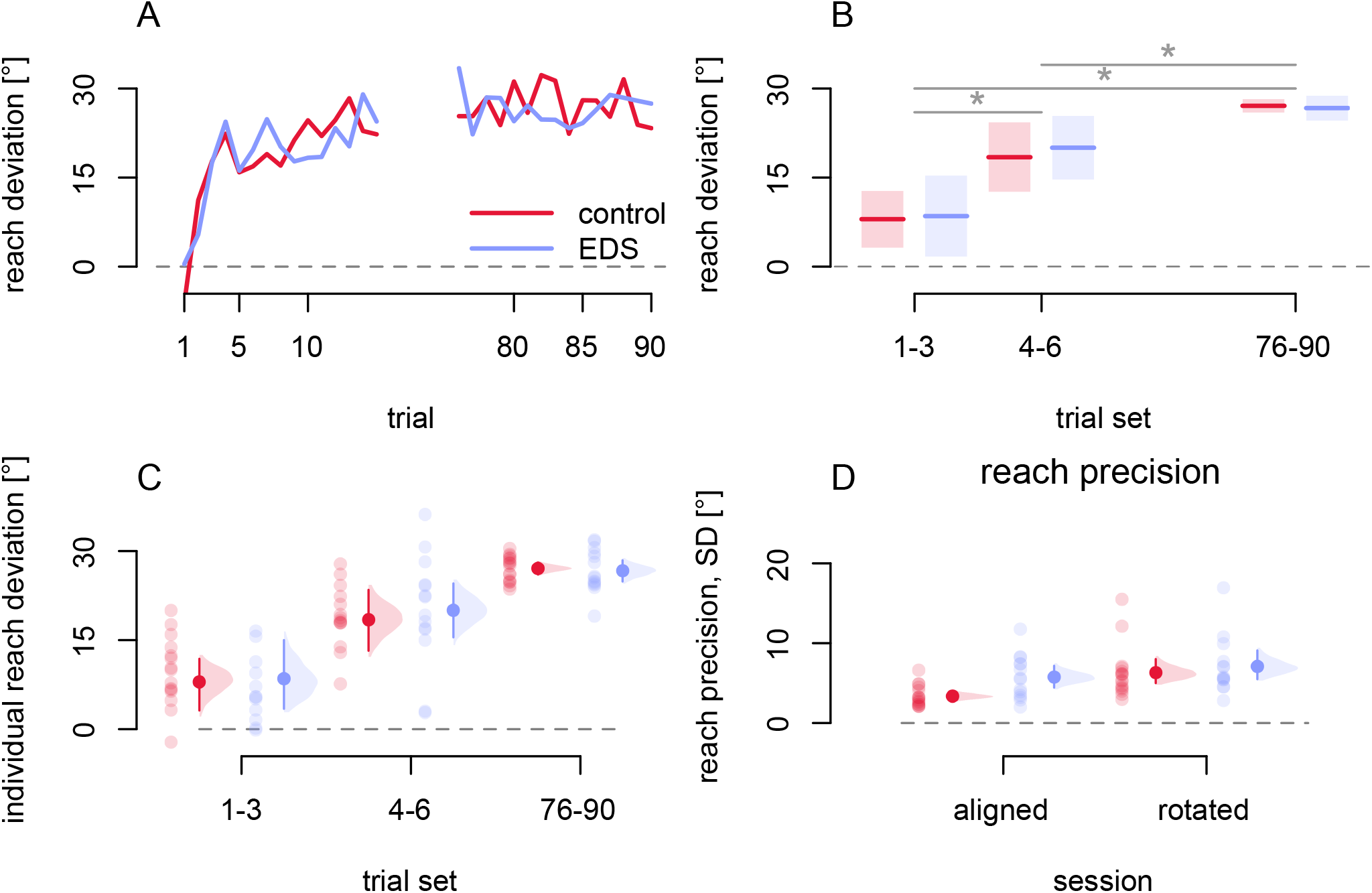
Rate of learning during adaptation. Controls are shown in red and EDS participants are shown in blue. Significant differences are indicated with an asterisk. Grey dashed line at the 0° mark indicates where aligned reaches are directed. **A-C:** The first and last 15 trials of rotated-cursor training are shown (A) across trials and (B-C) averaged for 3 sets of trials. Reaches directed towards 30° would mean that the hand had fully deviated to counter the perturbation. Solid lines are means and shaded regions are showing 95% confidence intervals (A-B), while individual data are shown as lighter-coloured dots. **D:** The standard deviation of target-normalized reach errors, for the last 15 trials of training in both the aligned and rotated sessions for each participant in each group (lighter dots). C-D. Dark dots and error bars correspond to the group mean and bootstrapped 95% confidence intervals.

### Reach Aftereffects

To measure implicit learning, we compared no-cursor trials both before (aligned) and after (rotated) adaptation (figures 4A & 4B) using a 2 × 2 mixed design ANOVA with training (aligned or rotated) as a within-subject factor and group as a between-subject factor. We confirmed the presence of reach after-effects with a statistically significant main effect of training (F (1, 28) = 133.19, p < .001, η^2^ = 0.607). However, we did not find a statistically significant main effect of group (F (1, 28) = 0.271, p = .607, η^2^ = 0.006), nor a statistically significant interaction between group and training (F (1, 28) = 0.030, p = .863, η^2^ = 0.0003). This suggests that, as expected, there is no difference between the groups in implicit learning.

**Figure 4.**
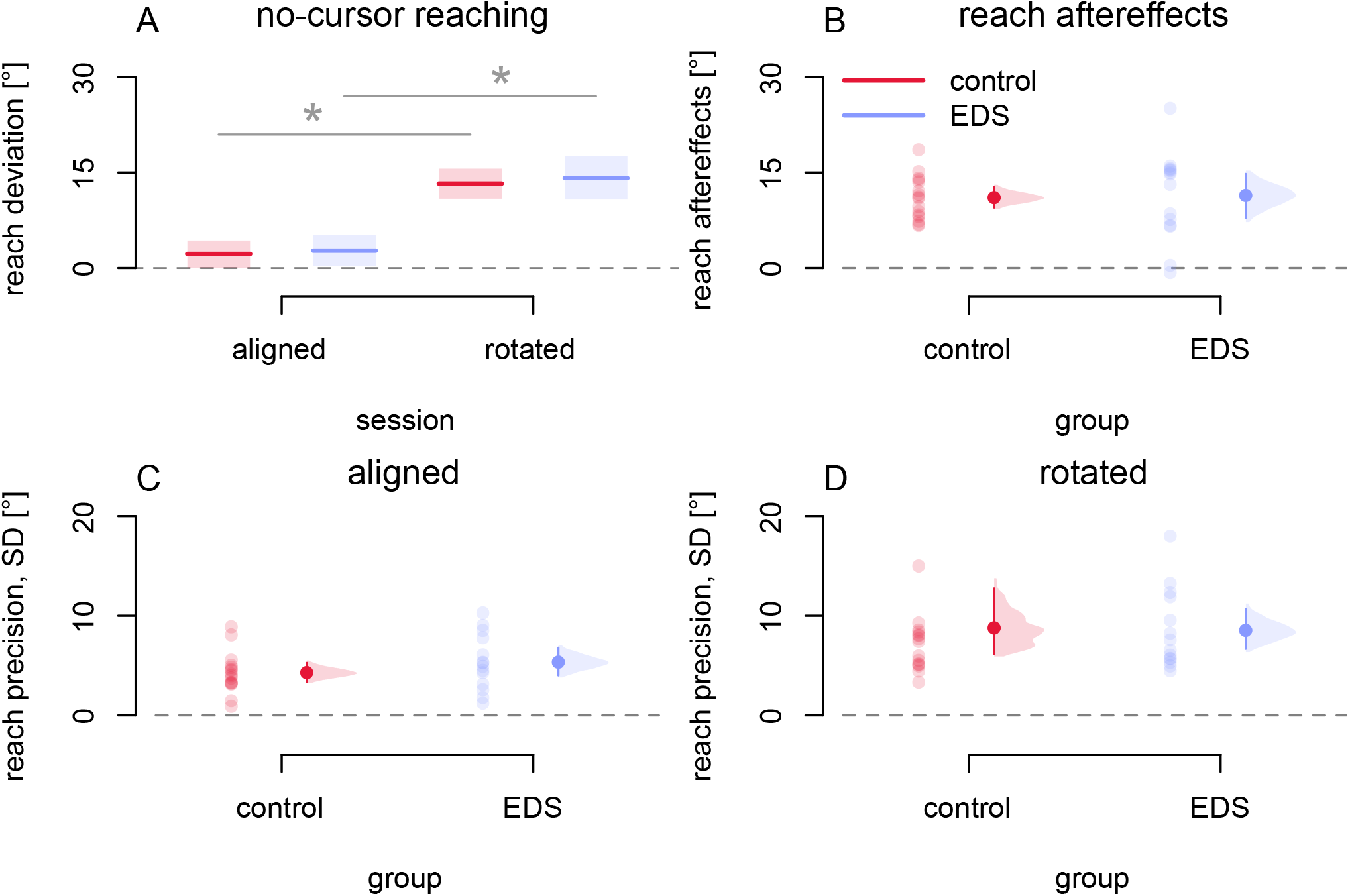
Measures of implicit learning. No-cursor reaches for controls are shown in red and those for EDS participants are shown in blue. Significant differences are indicated with an asterisk. The grey dashed line at the 0° mark indicates reaches that did not correct for the perturbation. **A:** Angular reach deviations of the hand per group before (aligned), or after (rotated), training. A reach deviation of 30° would indicate angular reach deviations equivalent to full compensation for the perturbation. Solid lines represent group means and shaded regions represent 95% confidence intervals. **B-D:** Individual participant data from each group are shown with transparent dots, while solid dots correspond to the group mean and bootstrapped 95% confidence intervals, respectively. **B:** Individual participant reach aftereffects (differences in reach deviations between the aligned and rotated sessions). **C-D:** Individual participant reach precision. The SD is calculated from all trials for each individual for the aligned session (C) and rotated session (D).

Closer to our main questions, to measure the precision of reaching when people cannot see their hand, we compared the standard deviation of target-normalized open-loop reach errors both before (aligned) and after (rotated) adaptation (figures 4C & 4D). We conducted a 2 × 2 mixed design ANOVA with training (aligned or rotated) as a within-subject factor and group as a between-subject factor. We found a significant effect of training (F (1, 28) = 10.747, p = .003, η^2^ = 0.163), such that reach scatter increased after rotated training. However, we found no significant effect of group (F (1, 28) = 0.119, p = .733, η^2^ = 0.002) nor any significant interaction between training and group (F (1, 28) = 0.303, p = .587, η^2^ = 0.005). A similar pattern was found for reaches made with a visible hand-cursor (figure 3D; refer to OSF for analyses). This suggests that EDS does not lead to greater variance in reaches.

### Hand Location Estimates

After confirming that EDS does not seem to affect baseline reaches, or adaptation, next we explored our main set of questions: whether EDS leads to differences in estimates of hand location, both before and after adaptation. Beginning with before adaptation (aligned session), to see if there are any differences in accuracy, or systematic errors, we plotted active localization biases (figure 5A; refer to OSF for analyses) and passive localization biases (figure 5B; refer to OSF for analyses) for hand positions ranging from 30° to 150°. As we can see in each of these figures, there is no such evidence of significant differences between the groups given that the means of these biases (and their 95% confidence intervals) overlap.

**Figure 5.**
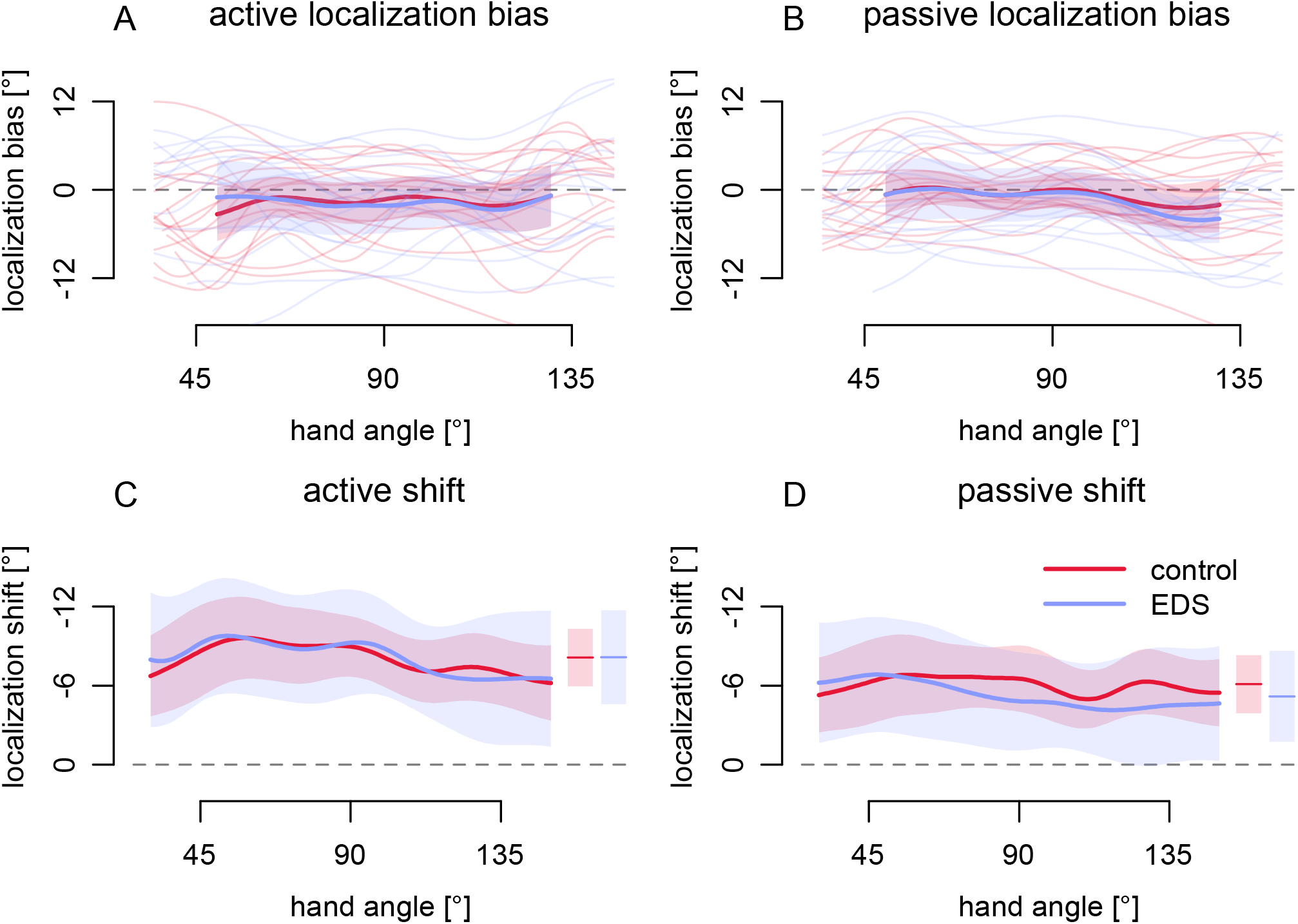
Hand localization estimates. Controls are shown in red and EDS participants are shown in blue. **A:** Active and **B:** Passive hand localization bias across the tested range during the aligned session. Bold, solid lines correspond to group means at each hand angle while shaded bars correspond to 95% confidence intervals for all panels. Individual participant data is shown with transparent lines. **C-D:** Visually-induced changes in hand location estimates for Active (C) and Passive (D) localization. Grey dashed lines at the 0° mark indicate the absence of shifts, while negative values indicate the direction of shifts consistent with the visual distortion. Horizontal lines correspond to group means collapsed across hand angles while shaded bars represent the 95% confidence intervals of these means.

Next, we investigated the effects of EDS on learning-induced ‘shifts’ in hand localization estimates (3^rd^ column of the last two tasks in table 1). We conducted a 2 × 2 × 2 ANOVA on localization error with group (EDS or control) as a between subject factor, as well as training (aligned or rotated) and localization type (active or passive) as within-subject factors. We found a significant main effect of training (F (1, 28) = 58.85, p < .001, η^2^ = 0.220), a significant main effect of localization type (F (1, 28) = 5.78, p = .023, η^2^ = 0.005) and a significant interaction between training and localization type (F (1, 28) = 17.77, p < .001, η^2^ = 0.009). This suggests that estimates of unseen hand location shifted following reach training with a rotated cursor and that the size of these shifts were slightly larger for active localization compared to passive localization (illustrated in figures 5C & 5D), as found in previous studies from our lab [10–14]. Since the focus of the current study is concerned with exploring these patterns in EDS, we then investigated group differences in active and passive localization shifts (by subtracting aligned localizations from rotated localizations to create a measure of localization shift) and conducted a 2 × 2 mixed design ANOVA on localization shifts with localization type (active or passive) as a within-subjects factor and group (EDS or control) as a between-subjects factor. We again found a significant effect of localization type (F (1, 28) = 17.769, p < .001, η^2^ = 0.059), that is active localization (5C) was slightly larger than passive localization (5D), and this difference can be seen when comparing figures 5C & 5D. However, we found no significant effect of group (F (1, 28) = 0.064, p = .803, η^2^ = 0.002) nor any significant interaction between localization type and group (F (1, 28) = 0.650, p = .427, η^2^ = 0.002). This suggests that there is no difference between the EDS and the control group in the magnitude of their localization shifts across either localization task.

### Localization Precision

Although, as expected, we did not find any group effects on any of the localization measures that reflect accuracy, we wanted to investigate the effect of EDS on precision of hand location estimates. We used standard deviations of hand localizations for each participant, then compared precision between groups (figure 6). We conducted a 2 × 2 × 2 mixed ANOVA with training (aligned vs rotated) and localization type (active vs passive) as within-subjects factors and group (control vs EDS) as a between-subjects factor. We found a significant main effect of group (F (1, 28) = 7.95, p = .009, η^2^ = 0.143), as well as a significant main effect of localization type (F (1, 28) = 13.21, p = .001, η^2^ = 0.050). However, we found no significant interaction between group and localization type (F (1, 28) = 0.92, p = .346, η^2^ = 0.004), which would suggest the significant group effect applies to all four conditions of hand localization. That is, the EDS group was less precise in all four conditions. However, in case the absence of a significant interaction was due to insufficient power, we decided to investigate this specifically using less conservative follow-up tests (one-tailed t-tests) to compare both groups across all 4 combinations of conditions (aligned active localizations, aligned passive localizations, rotated active localizations, and rotated passive localizations). We found no significant difference between groups for aligned active localizations (one-tailed t (23.69) = −1.25, p = .112, η^2^ = 0.055) (figures 6A & 6C). However, we confirmed the significant difference between groups in the other three conditions (Aligned Passive: one-tailed t (15.83) = −1.92, p = .037, η^2^ = 0.128; Rotated Active: one-tailed t (17.34) = −2.64, p = .008, η^2^ = 0.215; Rotated Passive: one-tailed t (26.02) = −2.48, p = .010, η^2^ = 0.183) (figures 6B & 6D). Consistent with the absence of a group & localization-type interaction, the results of these less-conservative, and uncorrected, t-tests suggest that precision in passive hand-localization was poorer for EDS compared to controls, as we predicted. That is, sense of proprioception is more variable in the EDS group when only afferent information is available. But when efferent information was also available, like in the active localization tasks, the difference was not so consistent, with the EDS group showing poorer precision following rotated training but not aligned training.

**Figure 6.**
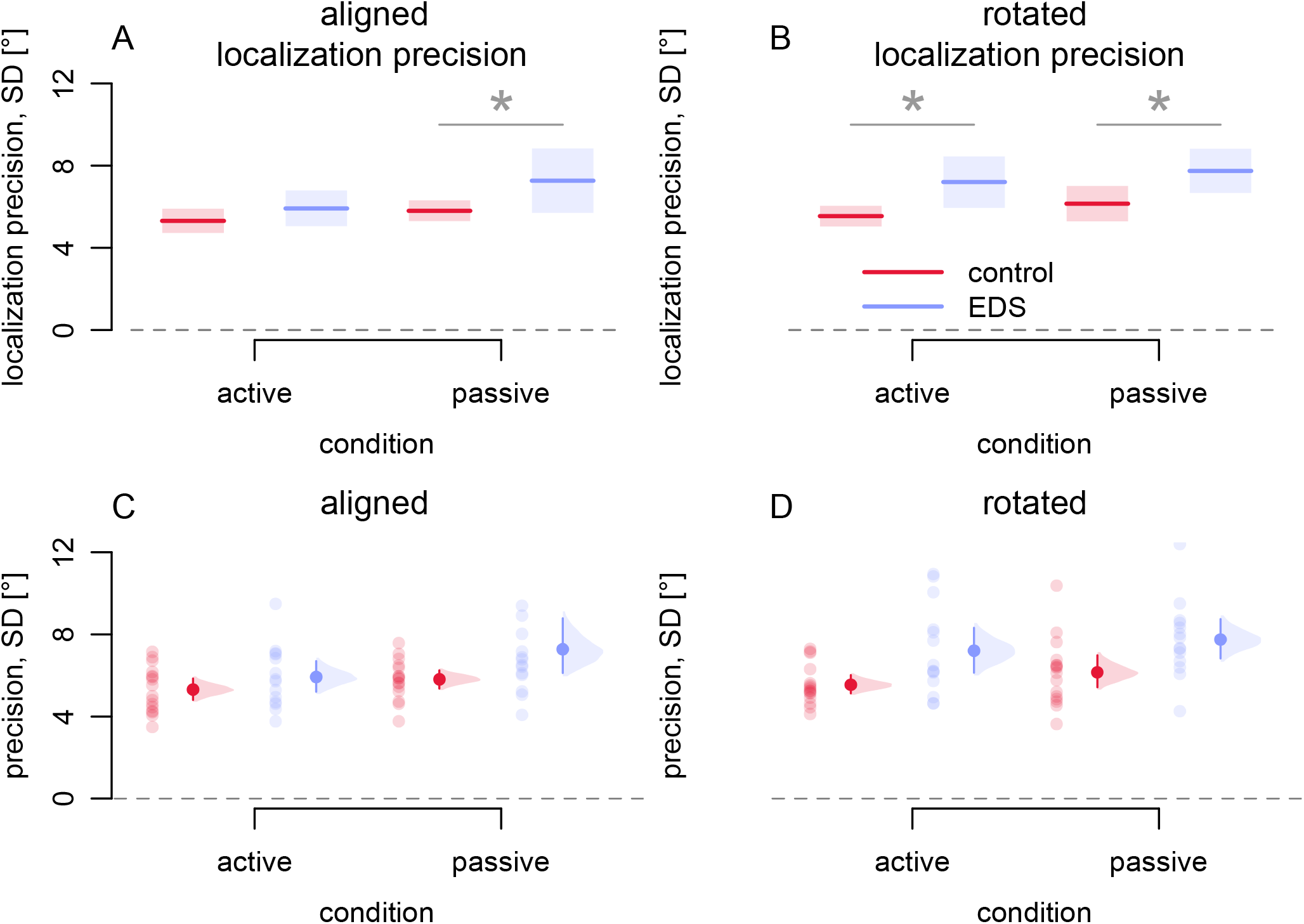
Localization precision. Controls are shown in red and EDS participants are shown in blue. Significant differences are indicated with asterisks. **A-B:** Solid lines represent group means and shaded regions correspond to 95% confidence intervals for hand localization precision (SD) in the aligned (A) and rotated (B) conditions, for both the active and passive hand localization tasks. **C-D:** Transparent dots indicate individual participant variability (SD) for hand localization in the aligned (C) and rotated (D) conditions for each group, and for both active and passive localization. Solid dots and error bars to the side of individual data correspond to group means and bootstrapped 95% confidence intervals.

Since variability in hand estimates is greater in the EDS group, then perhaps there is a relationship between joint hypermobility and the precision of limb localization. We used Beighton scores as a measure of joint hypermobility, measured in both groups, to investigate whether this is correlated with overall localization variance (variance was calculated across all 4 conditions for every participant to provide more power). Results of a Pearson correlation revealed that there was a significant relationship between joint hypermobility and measures of hand localization (figure 7; p = 0.036, r2 = 0.117). This suggests that those who are the most hypermobile tend to have the least precise (most variable) proprioceptive estimates of hand position.

**Figure 7.**
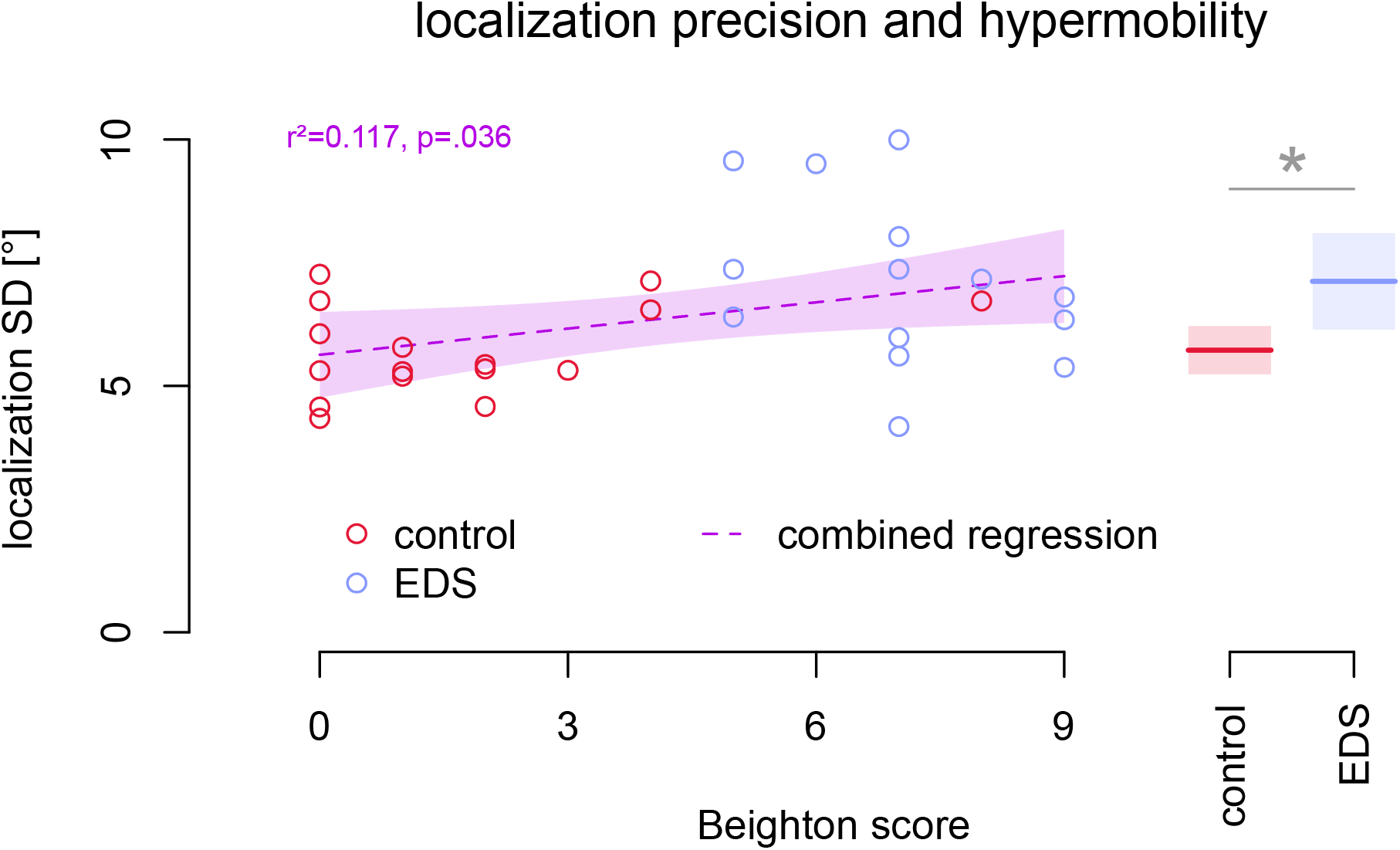
The relationship between localization precision and joint hypermobility. Localization standard deviations are plotted as a function of Beighton scores for EDS participants (blue) and controls (red). The dashed line corresponds to a regression line, while the purple shaded regions correspond to 95% confidence intervals. Solid lines represent group means, collapsed across all 4 localization conditions, and shaded bars correspond to 95% confidence intervals. The significant difference between the groups is indicated with an asterisk.

## Discussion

The goal of this study was to explore the effect of EDS on acuity and plasticity of hand proprioception. Specifically, our main goal was to quantify proprioceptive acuity of the upper arm, when only proprioceptive feedback was available (passive localization) or when afferent and efferent signals were both available (active localization). Our second goal was to better understand proprioceptive plasticity in EDS by having participants adapt their movements to a visual perturbation and then comparing shifts in both active and passive estimates of hand location across patients and controls. We found that people with EDS moved, and adapted their movements, just as well as controls. Accuracy and precision in reaching movements, and adaptation rates, did not differ between the two groups. Moreover, the accuracy by which the two groups estimated their unseen hand did not differ. The only aspect where people with EDS differed from controls was the precision of hand estimates, which was poorer for EDS patients when estimates were based on proprioception. When efferent information was also available, this difference disappeared. However, visuomotor adaptation disturbed estimates of hand position enough to produce poorer precision in estimates for EDS patients compared to controls for both types of hand localization. In summary, our results support the notion that proprioceptive sensitivity is different in EDS, and that poor proprioception can be overcome by additional efferent information.

The fact that we found no differences between patients and controls in any of our other measures in both the current study, and in a previous study [5], suggests that we have found an impairment that relates purely to proprioception. However, the current study suggests that efferent information may be sufficient to overcome poorer proprioceptive sensitivity. This would in turn explain why the poorer proprioceptive sensitivity did not lead to poorer reaches, or poorer reach adaptation, in this and our previous studies. Thus, arm-motor control and visuomotor integration processes in EDS are no different than in healthy controls. Given the pathophysiology of EDS, it is possible that the small deficit we have observed in this study could have both central and peripheral origins which are elaborated upon below.

Our current findings are in line with our past research, and the research of others, which suggests that proprioceptive precision of the upper limb is different in EDS. In our previous study [4], we also found that people with EDS showed twice as much scatter, compared to controls, when indicating the felt position of their unseen left hand. In the current study, we again show EDS did not differ from controls in the accuracy of their proprioceptive estimates, but in precision. While Rombaut et al. [7] found that there were no significant differences between Hypermobile-EDS patients and controls in absolute angular errors during their shoulder joint reposition test (the target-hand was passively placed, but its position was indicated both actively, like in our study, but also passively at target angles of 45° and 75°), they did find larger variations in angular errors (standard deviations that were 27-65% higher) for the patients in 3 of their 4 conditions (all except the passive 75° reproduction). What is new in the current study is the finding that these differences disappeared when people actively displace their own target-hand. Visuomotor adaptation, however, increased the uncertainty of the unseen hand location in EDS patients such that both types of hand estimates were less precise when compared to controls. Thus, proprioceptive variability was greater for patients in three of four hand localization conditions, which suggests some differences in proprioceptive sensitivity, and that this phenomenon is likely afferent in nature.

Studies which measure proprioception of the knee in those with joint hypermobility tend to find more obvious impairments. Rombaut et al. [7] also measured proprioception of the knee at two different detection angles. While they found that patients showed larger absolute angular errors than controls, the standard deviations of these errors were also much larger for patients (around 30% greater for most conditions but were double for active reposition of the 30° angle). Their findings are like those of Sahin et al. [8] where absolute angular errors were twice as large in EDS patients, compared to controls, for a knee joint reposition task. It is possible that we see greater differences in proprioceptive acuity at the knee joint since it is more of a weight-bearing joint; therefore, the knee may be more prone to repetitive stress-induced injury, which could have effects on proprioceptive acuity [7]. Another possibility is that the upper limbs, especially the hands, are more precisely represented in the somatosensory cortex, which could make people better able to compensate with varying sensory input when estimating the position of their hand [17]. Given that poorer proprioceptive precision is found for lower limbs and upper limbs in EDS patients, this suggests that proprioception may be compromised in those with hypermobility throughout the entire body.

An important finding from our current study is that the magnitude of joint hypermobility, as measured by Beighton Scores, is significantly related to our measures of proprioceptive precision; those who were the most hypermobile tended to also be those who were the least precise. This is like what we found in our first study, where joint hypermobility was found to be significantly related to uncertainty of proprioceptive estimates, but only at locations eccentric to the body midline [5]. However, we did not find a relationship between joint hypermobility and precision of hand location estimates in our other previous study, where participants had to reach to the felt location of their left hand, even though the precision of these estimates was double that of controls. We assumed that was due to the challenging nature of the task used, such that even those with mild levels of joint hypermobility would be likely to show proprioceptive impairments [4]. Regardless, we have found evidence (in the current study, and in the first) which suggests that proprioceptive variability is, at least partially, related to a person’s magnitude of joint hypermobility.

Why proprioception may be less precise in EDS is not clear, although some explanations have been proposed. Many types of EDS are known to be due to mutations in the genes that code for collagen, which could ultimately interfere with structural, and functional, aspects of the Extracellular Matrix (ECM) of connective tissues [18]. There are various neuroreceptors which are thought to contribute to our sense of proprioception (see [19] for a review) and these are all surrounded by connective tissues in our joints, muscular tissue, or skin, which may be affected by EDS; it is possible that activation of these proprioceptors is altered due to the interactions they would ultimately have with the ECM, but this possibility has never been explored. It is also possible that proprioceptive sensitivity in EDS is due to peripheral nerve damage, as small fibre neuropathy [20] and ulnar nerve subluxation/luxation [21] were found in those with EDS. Unfortunately, no studies to date have directly linked peripheral nerve damage to deficits in proprioception in this population. Recently, van Meulenbroek et al. [22] suggested that the proprioceptive differences commonly seen in adolescents with Hypermobile EDS could be due to physical deconditioning, because of kinesiophobia, since those with joint hypermobility are more prone to injuries and experience pain more intensely than the general population. Unfortunately, there have been no EDS proprioceptive studies to date which have included a measure of physical activity in their protocol, although a few studies have already found various forms of exercise to be effective in relieving pain, and sometimes proprioceptive deficits, in those with joint hypermobility [3, 8, 23]. Wearing compressive garments was also found to somewhat improve postural deficits in EDS [24], which researchers attributed to altered cortical representations of body schema that could be reorganized by enhancing cutaneous sensation. Given that many types of EDS often present with cutaneous abnormalities [2] and cutaneous receptors contribute to our sense of proprioception [19], it is possible that these are factors in the current study as well. It has been found that people with chronic low-back pain were significantly less accurate in their judgements of trunk rotations during a motor imagery task [25], although the possibility of disrupted, cortically held, representations of body schema in EDS has never been directly tested. It is our impression that proprioceptive deficits can arise for multiple reasons, but further studies need to be done to fully understand the impact of each of these potential causes on proprioceptive sensitivity in EDS.

### Limitations

We recognize that this study was likely underpowered. We determined that we would need 36 participants in each group in order to achieve 80% power. Unfortunately, we were not able to recruit as many participants as we hoped since EDS patients are rare and some people who were recruited were unable to attend due to health reasons. Then the COVID-19 pandemic occurred, which made it impossible to collect any more data. While our findings need replication, they should still be relevant to researchers in various fields.

Given the low power in our study, along with the a priori hypotheses we set out to explore, which were based on our previous findings showing poorer proprioceptive sensitivity in EDS, we opted to follow up our significant main effect of group for localization precision, but absence of a significant interaction between group and localization type, for each of the four localization conditions, using uncorrected one-tailed t-tests. Thus, we used less conservative follow-up tests to verify a non-significant interaction that would normally be interpreted to mean that the group difference applied to all the conditions. Nonetheless, we encourage readers to interpret the results more cautiously, given the low power, but our main reason for choosing these analyses was to reduce the likelihood of false negatives. We have included effect sizes for all our measures to aid readers in making their own interpretations.

Although we did have some a priori hypotheses, we did not pre-register our research protocol, or analysis with the OSF. However, all our data and analysis scripts are available on our OSF repository (see below).

## Acknowledgements

We would like to thank EDS Canada for promoting awareness of this project and allowing us to recruit participants through their support group. We would also like to thank Ayça Erdem for helping with data collection, along with Shanaa Modchalingam and Raphael Gastrock for their assistance with developing the paradigm. This work was supported by a NSERC Operating grant (DYPH).

## Data Availability

All data files are available from the Open Science Framework (https:llosf.iolt2jrsl).

